# Drug Repurposing: A Potential Therapeutic Strategy for the Treatment of Chikugunya Virus

**DOI:** 10.64898/2026.02.19.706773

**Authors:** Siphokazi S. Zondi, Letitia Shunmugam, Sphamandla E. Mtambo, Ndumiso M Buthelezi, Ayanda M. Magwenyane, Hezekiel M. Kumalo

**Affiliations:** Discipline of Medical Biochemistry, School of Laboratory Medicine and Medical Science, University of KwaZulu Natal, Durban South Africa

## Abstract

Chikungunya virus (CHIKV) infection is one of the major public health concerns in several countries around the world. CHIKV non-structural protein 2 (nsP2) is a promising drug design target due to the enzymes multifunctional properties that facilities viral replication and propagation. To date, there is an evident lack of preventative and therapeutic developments that can be used against CHIKV. Drug repurposing is a time saving and cost-effective method used for the development of new drugs. In this study, drug repurposing was implemented with the use of HIV/HCV protease inhibitors to inhibit the active site of nsP2. Molecular dynamics simulations and analysis revealed the stability of two drugs, Indinavir and Paritaprevir. Indinavir forms a hydrogen bond with a major residue, which closes the flexible loop, situated in close proximity to the active site. This conformational shift in the orientation of the enzyme prevents accessibility to the active site thus disrupting the nsP2 protein from functioning effectively in viral replication. In conclusion, the findings of this study identified Indinavir was identified as a promising CHIKV nsP2 inhibitor. This study will provide the basis to further facilitate the drug repurposing strategy as an alternative approach for drug design of CHIKV inhibitors as well as other viral families.

## 2. Introduction

Chikungunya is a viral disease that belongs to the Semliki Forest virus (SFV) serological complex within the Togaviridae family of the *Alphavirus* genus ^1^. This virus is primarily transmitted to human hosts through the bite of *Aedes aegypti* and *Aedes albopictus* mosquitoes. Chikungunya virus (CHIKV) is the primarily responsible for CHIKV infection in humans ^2^. Catergorised as an aphavirus, CHIKV was first identified in 1952, within the south region of Tanzania ^3^. Severe manifestations of CHIKV infection may include myocarditis, hepatitis, and neurologic complications such as encephalitis. In humans, there are several manifestations caused by CHIKV such as pyrexia, headaches, skin rash, myalgia, disabling arthralgia, gastrointestinal discomfort and conjunctivitis ^4,5^. Outbreaks of CHIKV have occurred in several countries across the Indian and Pacific Oceans as well as Europe, Asia, Africa and in the Caribbean. The risk of CHIKV being distributed throughout new geographical locations have been greatly catalysed by infected travelers. To date, there is no drug or effective vaccine that acts against the CHIKV infection ^6^.

Classified as a positive sense single-stranded RNA virus, CHIKV is rapidly evolving as a consequence of exposure to favourable climatic conditions that has also expedited geographical dissemination of the virus. The CHIKV genome is approximately 11.8 kilobase (kb) and encodes four non-structural (ns) proteins 1, 2, 3 and 4 (nsP1. nsP2, nsP3 and nsP4) and five structural proteins: capsid (C), envelope glycoproteins E1 and E2, as well as two small peptides E3 and 6K ^7,8^. The non-structural proteins have several function including host/virus interactions such as evasion of antiviral responses ^9^. Non-structural proteins are assigned specific functions that facilitate the process of viral replication and overall progation. The nsP1 is responsible for capping and methylation of newly formed viral genomic and sub genomic RNAs, nsP2 modulates viral replication, nsP3 serves as a functional component of the replication unit and nsP4 is an RNA-dependent RNA polymerase ^9–11^.

Amongst the non-structral proteins, nsP2 is the longest, approximately 798 amino acid residues in CHIKV ^9^. The nsP2 protein is a multifaceted enzyme, its proteolytic activity plays a crucial role in non-structural polyprotein cleavage, which is essential for viral replication. This functional mechanism is linked to thiol group deprotonation of cysteine within the active site by assistance of an adjacent histidine residue ^12,13^. The nsP2 protein contains a N-terminal RNA helicase domain as well as a C-terminal protease domain ^10^. The latter is associated with “cysteine protease activities and is also reffered to as thiol protease. In the N-terminal domain, there are functions of RNA-dependent 5’-triphosphatase, helicase and nucleoside triphosphatase ^12^. In its free form, nsP2 is responsible for transcriptional shut-off and induces cytotoxicity ^12^. Due to its substantially influential role in viral replication, nsP2 has been regarded as the most attractive drug design target by both the medicinal chemistry and pharmaceutical industry. This protein may hold the key for future developments of potently effective therapeutic agents and vaccines that act against alphaviruses ^12,14^.

The crystal structure of nsP2 has revealed the presence of a catalytic dyad made up of cysteine 10 (Cys10) and histadine 79 (Hie79). The Cys10 amino acid is present near the first helix of the N-terminal domain and Hie79 at a turn around β1 and β2 strands ^10,13^. The nsP2 contains a flexible loop near the active site and this loop contains an asparagine residue that blocks entrance to the substrate and if this residue is substituted to alanine, it reduced protease activity by a significant amount ^15^.

As a consequence of changing patterns involving vector distribution, escalating abundance in response to favourable climate change and an increase in vector-human interactions, CHIKV has been deemed a potential worldwide public health burden ^16^. Currently, there is no vaccine or sufficiently effective antiviral treatment that can be used to prevent or treat CHIKV infection. Due to the severity of untreated CHIKV infection, other complications may develop such as arthritis, neurological disorders and even death in some cases ^13^. Treatment has been limited to those that can alleviate symptoms, such as antipyretics, analgesics, corticosteroids and non-steroidal anti-inflammatory drugs (NSAIDs) ^17^. For this reason, it is highly important to continue antiviral research in effort to discover potential therapeutic compounds as well as possible preventative approaches such as vaccines that are directed against CHIKV. A promising approach in drug discovery and design known as “drug repurposing” has been brought forward as a promising therapeutic strategy in antiviral studies.

The design and development of new drugs to acquire FDA approval is not only a costly process, but it also takes up a lot of time. Repurposing drugs that already exists means that the risks of these drugs, if any exist, are already known and that these drugs have been approved by regulatory agencies ^18^. Drug repurposing is an attractive and effective process when it comes to drug development for neglected tropical diseases. This is particularly true as there are challenges in the sense of low market value, even though there is an increased need for these drugs. Repurposing already existing FDA-approved drugs is not only a cost-effective and less time consuming stratergy but the deliberate implementation of naturally-sourced compounds can potentially provide alternatives to the development of new antiviral therapeutic approaches.

Curcumin, is a well-researched natural active compound that is derived from rhizome of turmeric (*Curcuma longa*). This compound has been used over the years in traditional medicine to treat a wide variety of ailments including diabetes ^19^, cancer ^20^, sexually transmitted diseases ^21,22^, inflammatory-related disorders ^23^, bacterial, fungal and viral infections ^24,25^. Curcumin has been shown to be an antiviral compound with activity against complex viruses such as Dengue virus ^26^, herpes simplex virus ^21,22^ and HIV-1 ^22,27^. In 2017, Mounce and co-researchers investigated the effects of curcumin on CHIKV, the study revealed that the natural compound substantially reduced the activity of CHIKV thereby decreasing the overall infection rate if the virus ^24^. The exposure of CHIKV showed an IC50 of 3.89 µm curcumin. However, the safety index of the active compound was not satisfactory. Contrastingly, its derivative, demethoxycurcumin showed better safety profile and higher potency. This may be attributed to dmethoxycurcumin possessing lipophilic properties, thereby allowing the derivative compound to interfere with the viral membrane of CHIKV thus contributing to its antiviral mechanism of action ^10^.

In recent times, HIV protease inhibitors-nelfinavir and lopinavir have been successfully re-profiled as anticancer agents following drug repurposing investigations ^28,29^. These findings provided a fundamental basis for the medicinal chemistry and pharmaceutical industry that allowed the screening of previously FDA approved drugs thus utilising the drug repurposing approach on a wide range of inhibitors against a broad spectrum of viruses.

The development of peptidomimetic inhibitors targeting CHIKV nsP2 protein is considered a success in lead development targeting of this re-emerging virus. There are structural similarities between these peptidomimetic inhibitors and HIV/HCV protease inhibitors, which prompted the use of HIV/HCV protease inhibitors for CHIKV drug re-profiling ^4^.

In this study, the drug repurposing strategy was investigated with regard to CHIKV. Through extensive application of various molecular and bioinformatics tools, structural and dynamic properties of the free enzyme as well as several bound complexes, were expressed and compared. This study highlighted the importance and indispensability of drug repurposing in antiviral therapy. It is with great hope that this study will uncover already existing drug molecules as possible antiviral agents against CHIKV as well as a wide spectrum of other viruses.

## 3. Computational Methodology

### 3.1 System Preparations

The X-ray crystal structure of nsP2 of the CHIKV was extracted from the Protein Database (PDB code: 3TRK) (www.rcsb.org) ^10^. The various HIV/HCV protease inhibitor: Amprenavir, Atazanavir, Darunavir, Fosamprenavir, Indinavir, Lopinavir, Nelfinavir, Ritonavir, Saquinavir, Tipranavir, Asunaprevir, Boceprevir, Grazoprevir, Paritaprevir, Simeprevir and Telaprevir structures were constructed and optimized using SMILES strings available on UCSF Chimera software and Avogadro ^30^. The three dimensional structures of the receptor and the ligands were prepared using UCSF Chimera software ^31,32^.

### 3.2 Molecular Docking

Molecular docking was used to predict binding affinities and optimized conformations of the various ligands within the CHIKV nsP2 active site (grid box spacing of 0.375 Angstroms (Å) and x,y,z dimensions of 56, 74 and 42 Å). The docking software employed in this study was AutoDock Vina ^33,34^. The three top rated docked results with optimum poses and highest binding affinities to the CHIKV nsP2 binding site were selected and thereafter subjected to Molecular dynamic simulations. Identification of the binding site used for this study was previously confirmed by Kumar and co-investigators (2019) ^13^.

### 2.3 Molecular Dynamic (MD) simulation

Molecular Dynamic (MD) simulations is essential as it gives a focused outlook into the biomolecular process of biological systems. It is a very powerful tool that is used to gain in-depth knowledge of physical activities of molecules and atoms. The MD simulations were carried out using the Particle Mesh Ewald Molecular Dynamics (PMEMD) engine provided through GPU-accelerated Amber and ff14SB AMBER force field ^35–37^. The LEaP module was applied to solvate and neutralise the systems through addition of chloride, sodium and hydrogen atoms while being immersed in an orthorhombic box of TIP3P water molecules, all atoms were within 10 Å of the box parameters. A further 1000 steps of unrestrained conjugate gradient minimization were performed on all systems. Harmonic restraints of 10 kcal/mol Å^-2^ for solute atoms and Langevin thermostat with collision frequency of 1.0 ps^-1^ were used in order to heat all systems from 0 to 300 Kelvin (K) for 50 picoseconds (ps) in the canonical ensemble (NVT). All systems were equilibrated at 300 K for 500 ps and a constant pressure of 1 bar which was maintained by a Berendsen barostat. This was done to resemble an isobaric-isothermal ensemble (NPT). The SHAKE algorithm was used for the constraint of hydrogen bonds. The SPFP precision model was used for all simulations at a 2 (femtosecond) fs time step ^38,39^.

### 2.4 Post Dynamic Analysis

MD simulations were run continuously for 100 nanoseconds (ns) at a constant pressure of 1 bar and a constant temperature of 300 K. The coordinates of all the systems were saved individually every 1 ps and the trajectories were assessed using the CPPTRAJ and the PTRAJ modules in the Amber14 package ^38^. Post dynamic analysis that was performed includes; root mean square deviation (RMSD), root mean square fluctuations (RMSF), radius of gyration (RoG), dynamic cross correlation (DCC) and principal component analysis (PCA). Graphical and visualization software programs were extensively used throughout this study, including UCSF Chimera ^32^ to visualize projected trajectories and Origin software (OriginLab, Northampton, MA) for the construction of graphs.

#### 2.4.1 Root mean square fluctuations

Root mean square fluctuations (RMSF) is the fluctuation of individual or a group of atoms relative to their average position during a molecular dynamics (MD) simulation ^38–40^. This analysis gives insight into the flexibility of residue regions of the apo and bound systems of 3TRK. It is calculated as follows:

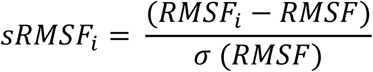

where RMSF_i_ is the RMSF of the i^th^ residue from which the average RMSf is subtracted. It is then divided by the standard deviation of RMSF [σ(RMSF)] to give the standardized RMSF [sRMSF_i_].

#### 2.4.2 Radius of gyration

The radius of gyration (RoG) is used to describe the root mean square deviation (RMSD) of all the atoms from the centre of gravity of an enzyme molecule. This analysis looks at the compactness of a protein system during MD simulations. The RoG is estimated using the following equation:

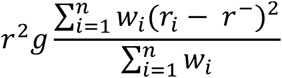

where r_i_ is the position of the i^th^ atom and the r is the centre mass of atom i. The mean value is calculated by taking the RoG values over the number of frames in a trajectory ^38,41^.

#### 2.4.3 Dynamic cross correlation

Dynamic cross correlation is used to determine the quantification of the correlation coefficients of motions between the atoms of a protein. The residue-based fluctuations of the system during the MD simulation were determined using the CPPTRAJ module. The following equation represents the calculation of DCC:

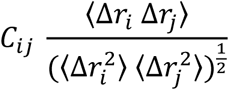

where C_ij_ is the cross correlation coefficient and varies from -1 to +1. A fully correlated motion represents the upper limits during the simulation and the lower limits are represented by an anti-correlated motion. The term i represents the i^th^ residue and j the j^th^ residue. Δr_i_ and Δr_j_ represent the displacement vectors for the i^th^ and j^th^ residues, respectively.

#### 2.4.4 Principal Component Analysis

Principal Component Analysis (PCA) is also known as essential dynamics (ED) and is an extremely versatile tool used for trajectory analysis. Principal Component Analysis has been seen to be powerful and robust and it also opens new opportunities in visualizing and exploring the protein cavity dynamics. It describes the eigenvectors and eigenvalues, which indicates the direction of motions and amplitudes of the protein, respectively (Cocco et al., 2013). The PCA was performed on C-α atoms over 1000 snapshots, at a time interval of 100 ps. The CPPTRAJ module in AMBER 14 was used to compute the first two principal components, PCA1 and PCA2. The corresponding PCA scatter plots were generated using Origin software (OriginLab, Northampton, MA).

#### 2.4.5 Thermodynamic Binding Free Energy Calculations

The binding free energy (BFE) calculation includes both entropic and enthalpic contributions and it is a crucial method used to observe the detailed binding mechanism between a protein and a ligand ^38,39^. This method provides valuable insight into the protein-ligand association of a complex and this is taken as the endpoint energy calculation. The binding free energy calculations for the systems were computed using the Molecular Mechanics/Generalized Born Surface Area (MM/GBSA) approach ^42,43^. In this study, the binding free energies were averaged over 1 000 snapshots that were extrapolated from the 100 ns trajectory. The explicit solvent used in the MD simulation was removed and substituted with a dielectric continuum as per MM/GBSA protocol. The changes in each term between the apo and bound systems were calculated and contributed to the total BFE. Molecular mechanics force fields were then used to calculate the energy contributions from the coordinates of the atoms of the ligand, enzyme and complexes in a gaseous phase ^38^. The equations below explain the process in which the binding free energies (*ΔG*) were assessed:

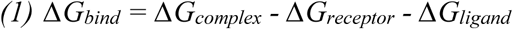

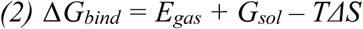

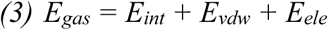

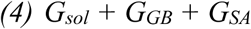

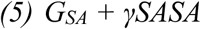

Where *E_gas_* is the gas-phase energy, *E_int_* the internal energy, *E_ele_* the electrostatic energy and *E_vdw_* is the van der Waals energy. The gas-phase energy, *E_gas_*, is measured directly from the force field terms, FF14SB. The solvation energy, *G_sol_*, is calculated by summing up the contributions from the polar and non-polar states. By solving the GB equation, we are able to determine the polar solvation contribution, *G_GB_*. The non-polar solvation contribution, *G_SA_*, is determined from SASA (solvent accessible surface area) using a water probe radius of 1.4 Å. T represents the temperature and S the total solute entropy. The solute and solvent dielectric constants are set to 1 and 80, respectively ^38^.

## 4. Results and Discussion

From the table, Paritaprevir, Grazoprevir and Indinavir were accompanied by the highest binding affinities following molecular docking. Thereafter, apo, Paritaprevir, Grazoprevir, Indinavir and Demethoxycurcumin were individually bound to the CHIKV protein receptor and MD simulations were conducted over a time period of 100 nanoseconds (ns).

### 4.1 Stability of the apo and bound systems of 3TRK

The root of mean square deviation (RMSD) is used to identify the stability of the C-α backbone in the apo and bound systems. Structural instability indicates the increased mobility of the C-α backbone, and it is represented by a high RMSD value. The lower RMSD values indicate decreased backbone mobility and therefore a more stable system. The RMSD plot is shown in figure 3 for Demethoxycurcumin, Grazoprevir, Indinavir, Paritaprevir ligands complexed to the enzyme and the Apo enzyme.

**Figure 1:**
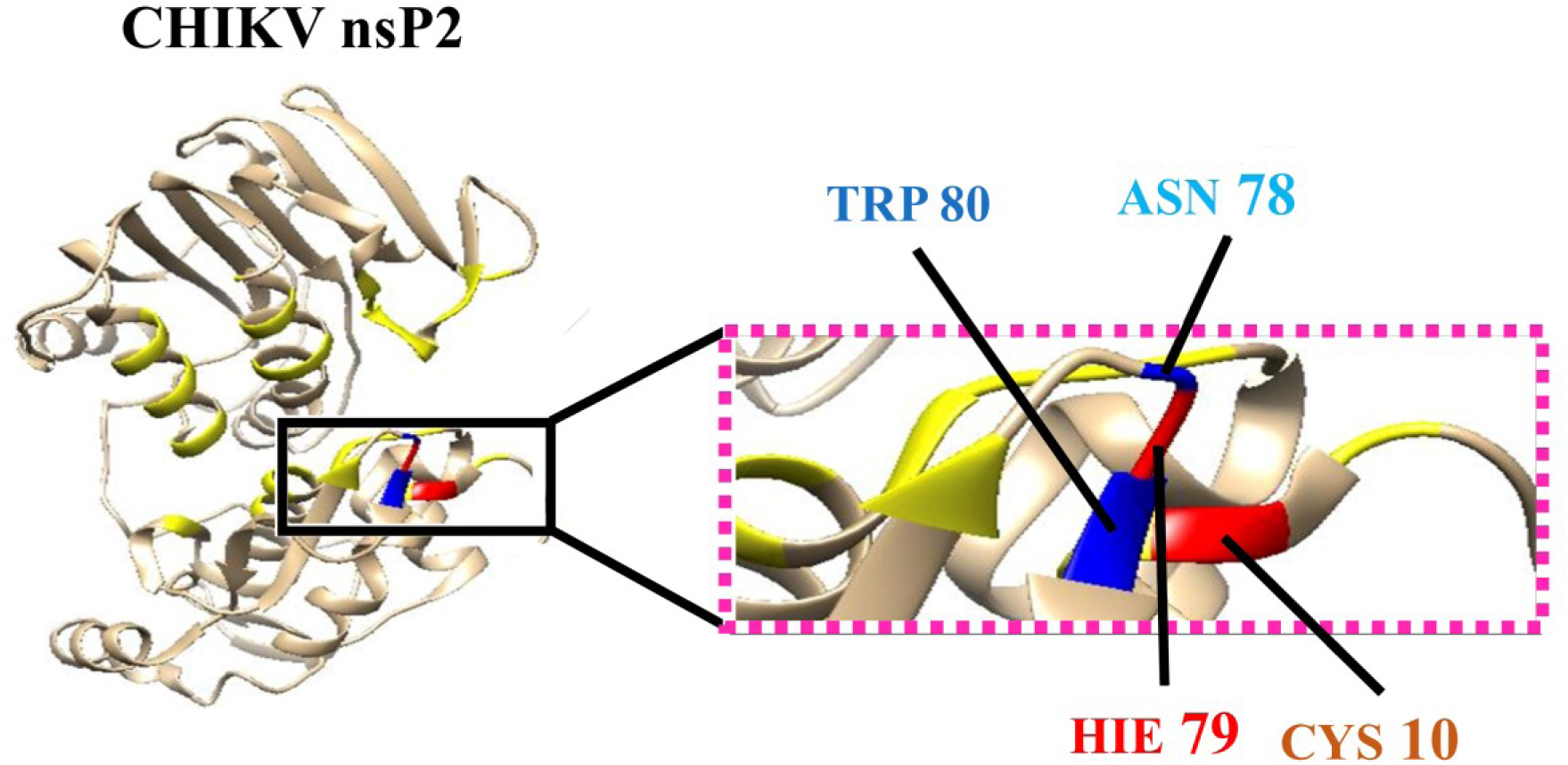
Crystal structure of nsP2 (PDB: 3TRK) showing the active site (yellow), the catalytic dyad Cys10 and Hie79 and the substrate binding residues Asn78 and Trp80.

**Figure 2:**
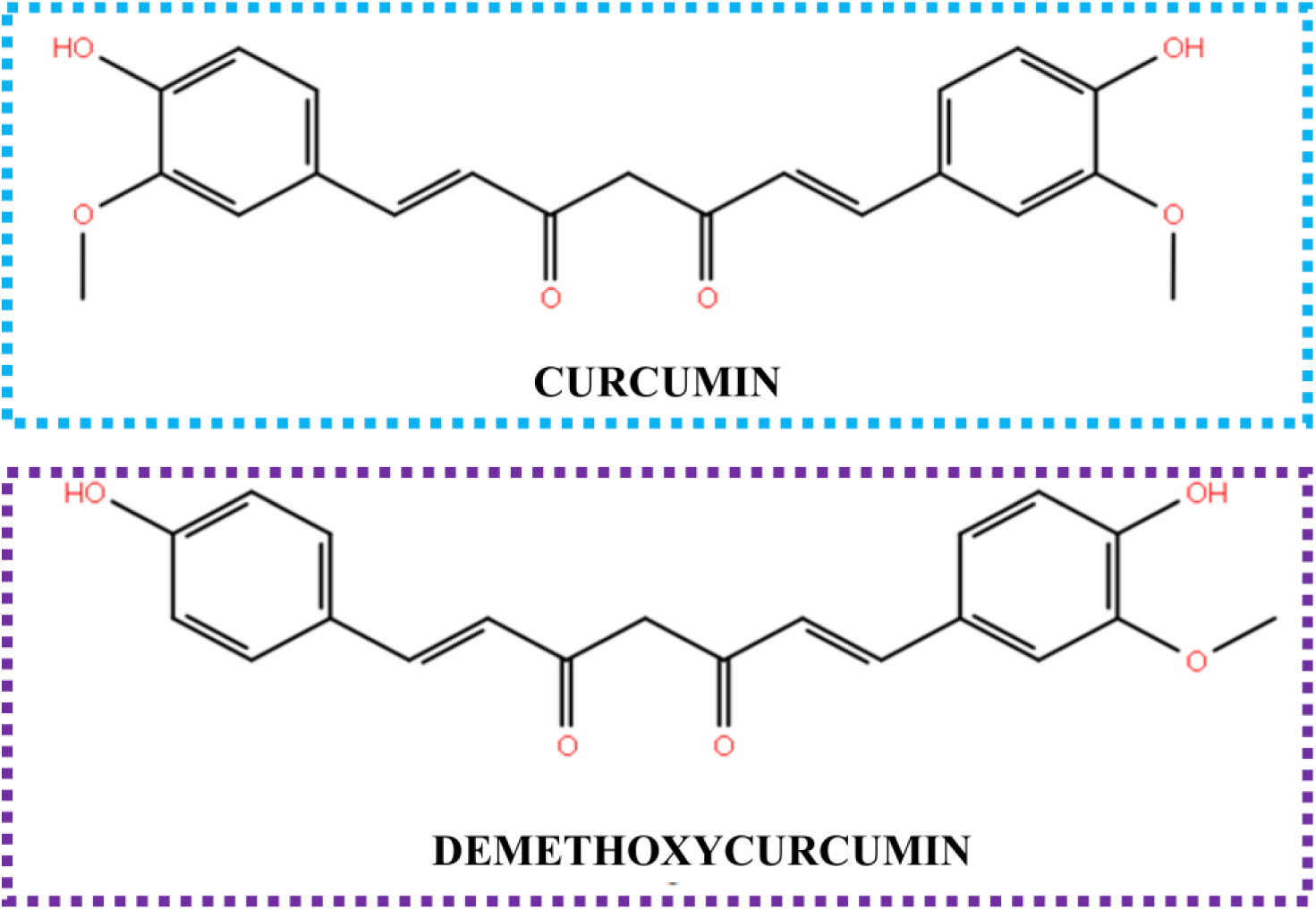
The 2-dimensional chemical structure of Curcumin and Demethoxycurcumin.

**Figure 3:**
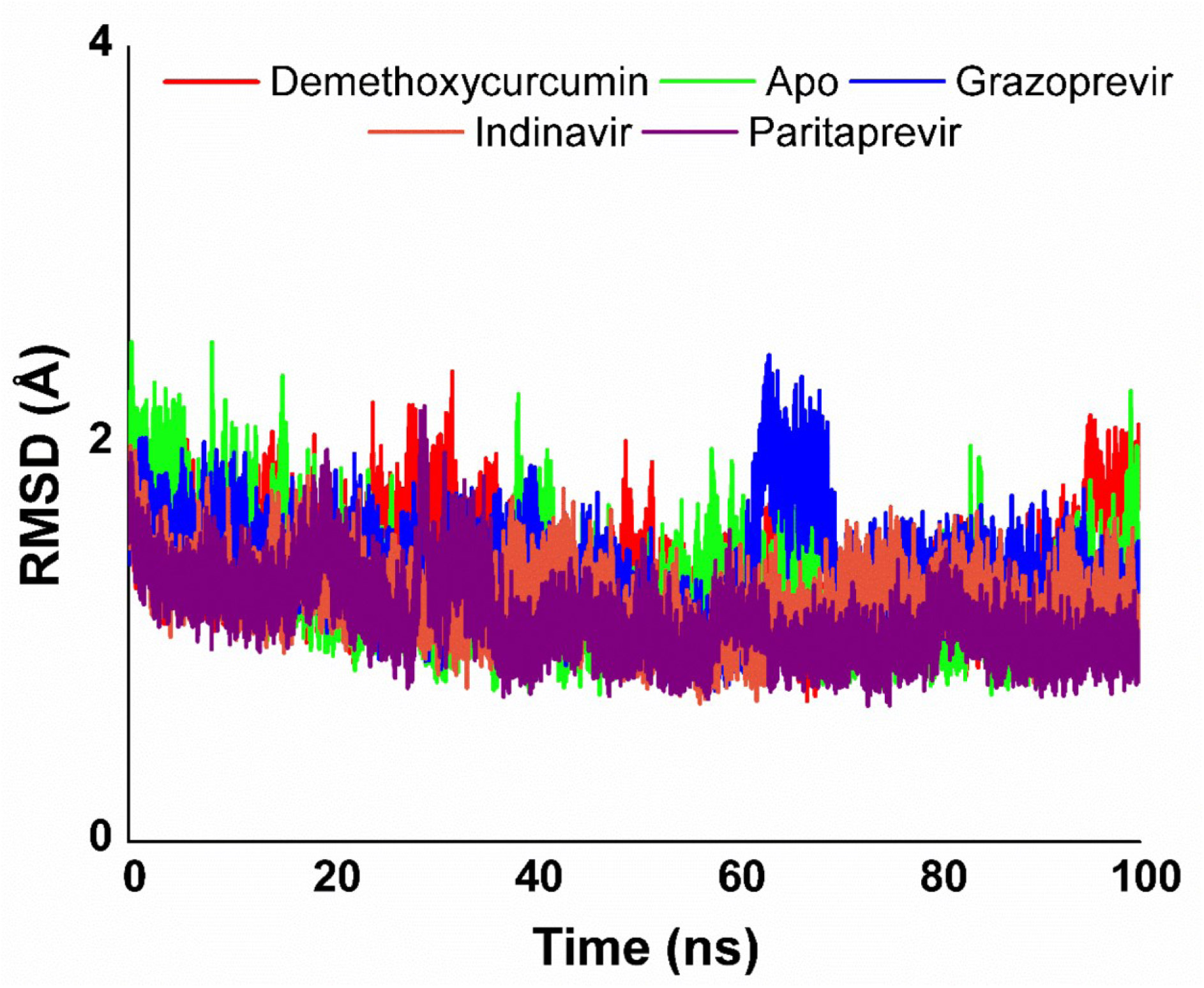
The RMSD of the free enzyme and complexes with Demethoxycurcumin, Grazoprevir, Indinavir and Paritaprevir were recorded over 100 ns MD simulations.

The RMSD graph above displays that all the systems were mostly stable throughout the simulation, except for Grazoprevir which showed some significant fluctuations. The major fluctuations of Grazoprevir were seen between 60 ns and 70 ns. From Figure 3, The most stable system was identified as Paritaprevir. Both, Indinavir and Paritaprevir demonstrated higher stabilities in comparison to Demethoxycurcumin. All systems had RMSD values less than 2 Å, which indicates that the equilibrium of all MD simulations is reliable ^44^.

### 3.2 Conformational Fluctuations of 3TRK

#### 3.2.1 Root Mean Square Fluctuation

Root mean square fluctuation (RMSF) is used to show the fluctuation of each residue during the 100 ns MD simulations. This gives insight into the regions of a protein that are flexible due to physiological changes that may have been caused by the binding of ligands to the enzyme. The RMSF of all residues were calculated for structure backbone flexibility and are shown in Figure 4.

**Figure 4:**
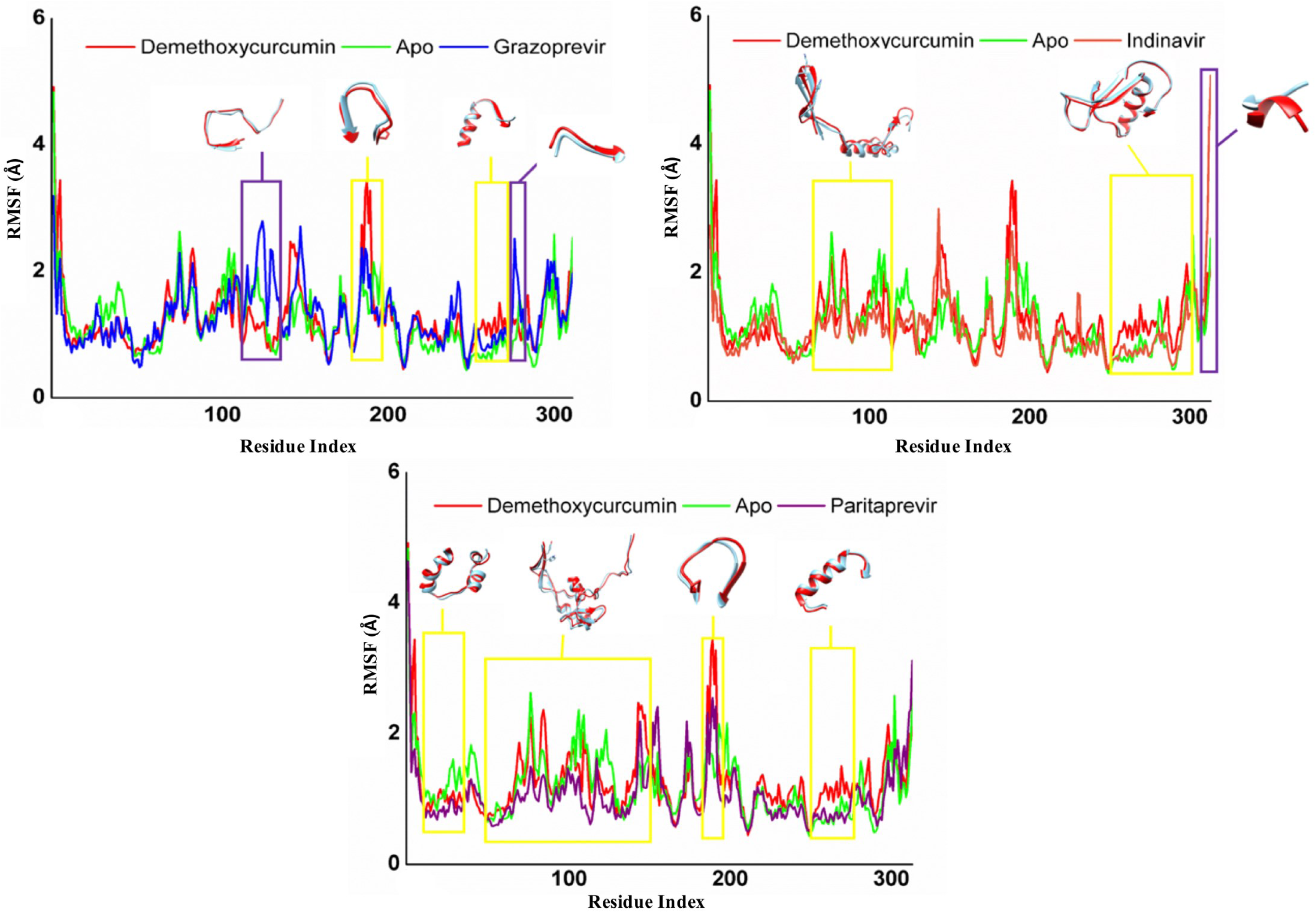
The RMSF of the Apo, Demethoxycurcumin, Grazoprevir, Indinavir and Paritaprevir.

Higher C-α atomic fluctuations values represent more flexible movements in relation to the average position of the residues during simulations. Lower C-α atomic fluctuations values symbolize the reduction of flexibility ^38^.

The analysis of the residue dynamic behaviour is significant as it provides in-depth knowledge into the type of bonding occurring between the ligand and enzyme residues. The graph of RMSF displays results that advocate to those of RMSD. The systems Grazoprevir, Demethoxycurcumin complexes and free enzyme displayed more flexibility in their residues with averages of 1.28416 ± 0.46934 Å, 1.26558 ± 0.5322 Å and 1.20642 ± 0.49075 Å, respectively. The most stable system was that of Paritaprevir followed by Indinavir with averages of 1.05994 ± 0.47159 Å and 1.11703 ± 0.48737 Å, respectively. In all the systems there are points where the Apo enzyme and the complex of Demethoxycurcumin show less fluctuations than the complexes of Grazoprevir, Indinavir and Paritaprevir. These points are circled in purple and indicate that the bonds formed between the enzyme and grazoprevir, indinavir and Paritaprevir caused fluctuations ^13^. The decrease in residual fluctuations of the Grazoprevir, Indinavir and Paritaprevir systems in comparison to the free enzyme and the Demethoxycurcumin complex, represents the formation of bonds with more van der Waals and electrostatic forces. The overall decrease in flexibility of the complexes means that the protein cannot carry out its function. The virus is not able to replicate efficiently after the binding of the ligands as the enzyme is not able to open and allow RNA to enter and exit, meaning that replication and propagation will not occur.

#### 3.2.2 Dynamic Cross Correlation

Dynamic cross correlation (DCC) analysis was performed in order to determine the dynamic correlations of the apo and bound systems. This method of analysis determines the position of the C-α atoms throughout the various simulations. The DCC matrix is represented in figure 5.

**Figure 5:**
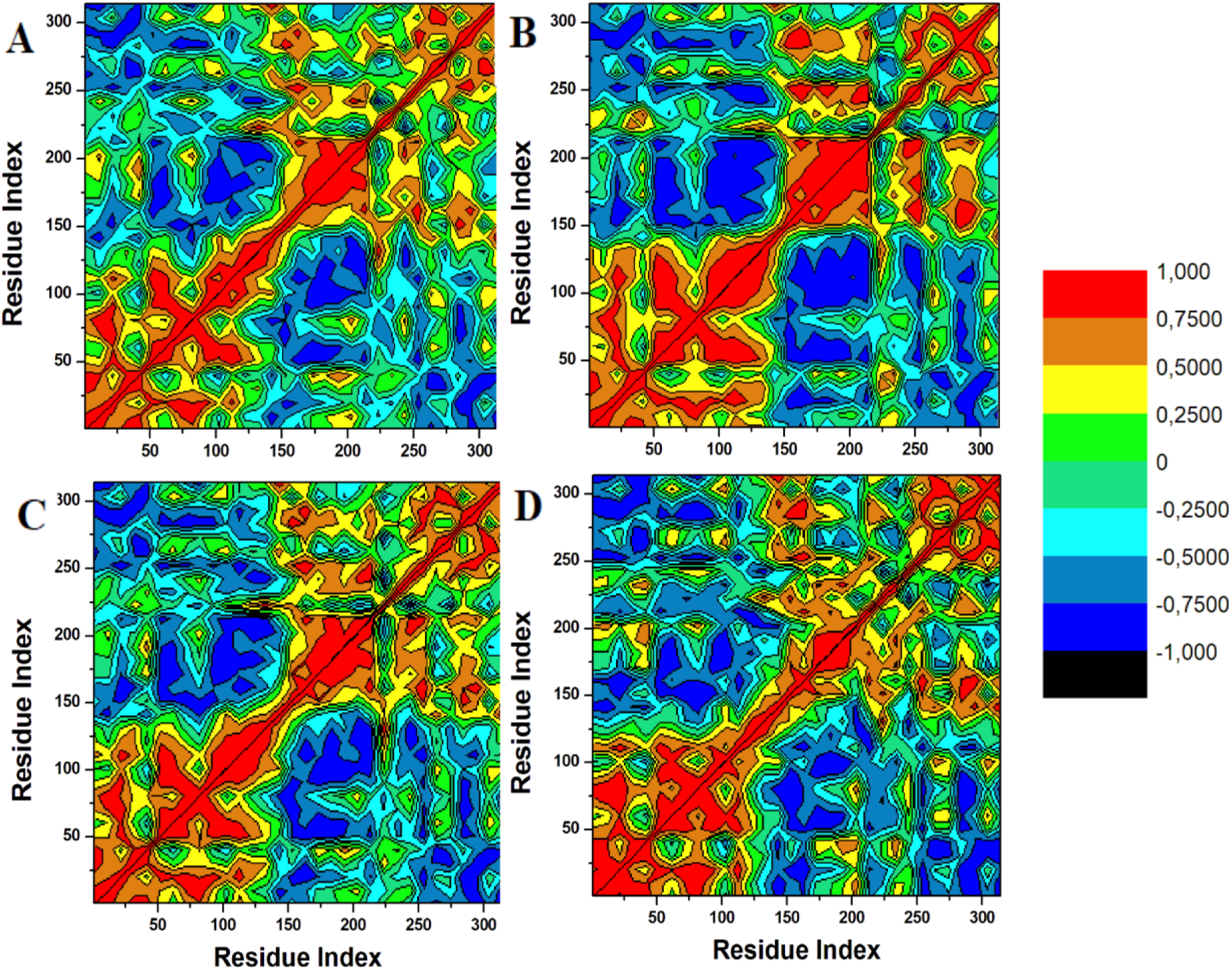
The dynamic cross correlation matrix of the complexes of (**A**) Demethoxycurcumin, (**B**) Grazoprevir, (**C**) Indinavir and (**D**) Paritaprevir.

The colours red to yellow represent highly positive correlated C-α atom movements and the colours blue to black represent negative or anticorrelated movements. The diagonal regions show obvious correlated movements. There is a similar pattern of correlated movements displayed by all systems, however, the unbound system shows higher levels of correlation. The system of the Grazoprevir complex displays an increase in correlated movements between the region of Arg 100 to Lys 150. These results support those observed in the RMSF where grazoprevir showed higher fluctuations in this region. There is a decrease in correlated movement around the region of residue 300 in the Demethoxycurcumin system. These results are in agreement with those seen in the RMSF, where Demethoxycurcumin showed to decrease the fluctuations of the residues in this region more than the other systems. The DCC results further prove that the formation of complexes of the enzyme 3TRK stabilizes the residues.

### 3.3 Radius of Gyration (RoG)

Radius of gyration (RoG) is used to show the compactness of the protein structure and gives insight into the complex stability of the various systems during the 100 ns MD simulations. Figure 6 shows the plot of Rg for the 5 systems.

**Figure 6:**
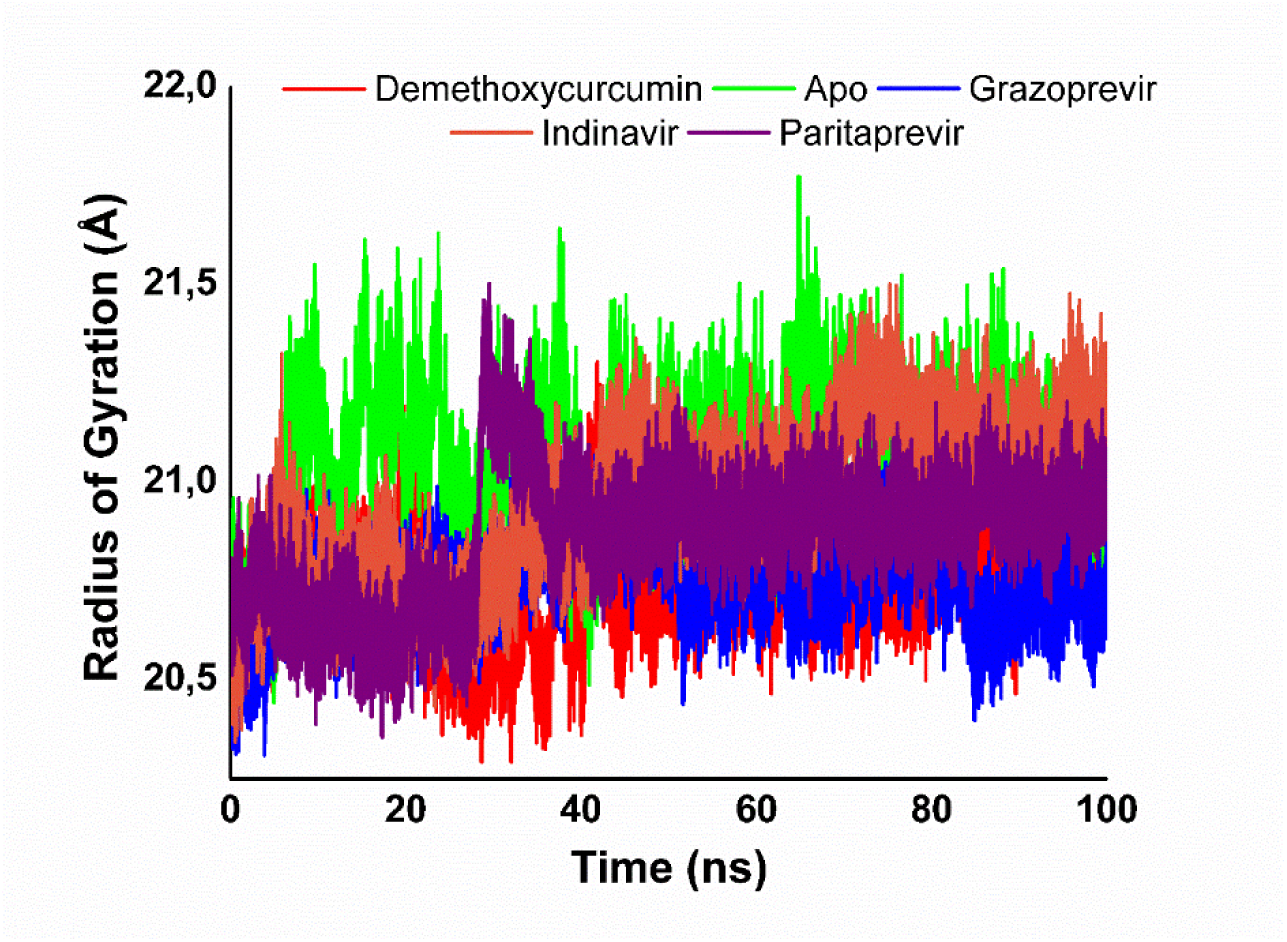
Radius of Gyration (RoG) of the Apo enzyme and Demethoxycurcumin, Grazoprevir, Indinavir and Paritaprevir complexes over 100 ns MD simulations.

**Figure 7:**
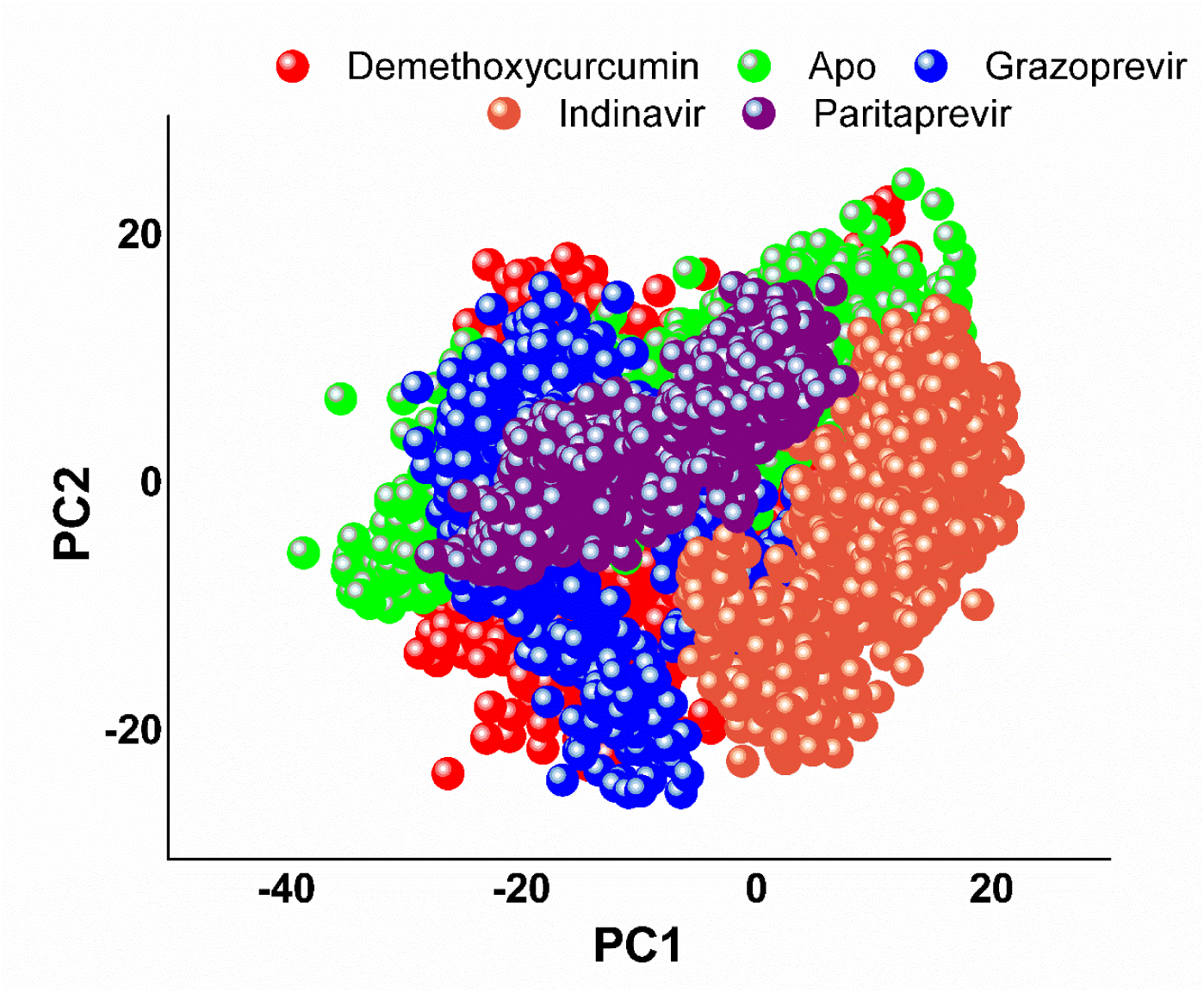
The PCA of the free enzyme of 3TRK and the complexes of 3TRK with Demethoxyxurcumin, Grazoprevir, Indinavir and Paritaprevir over 100 ns MD simulations.

**Figure 8:**
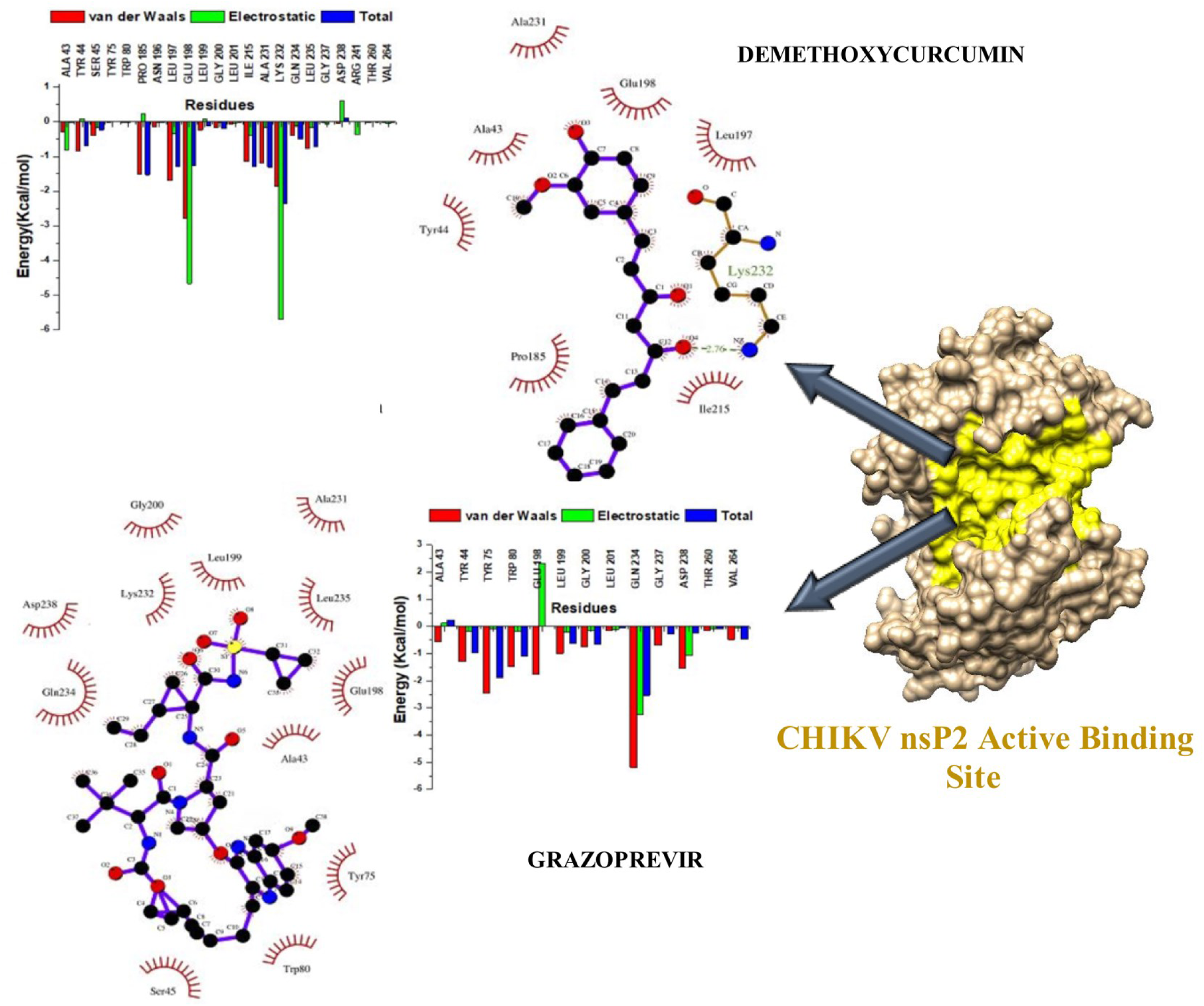
The per-residue free energy decomposition of Demethoxycurcumin and Grazoprevir bound to the active binding site of CHIKV nsP2.

**Figure 9:**
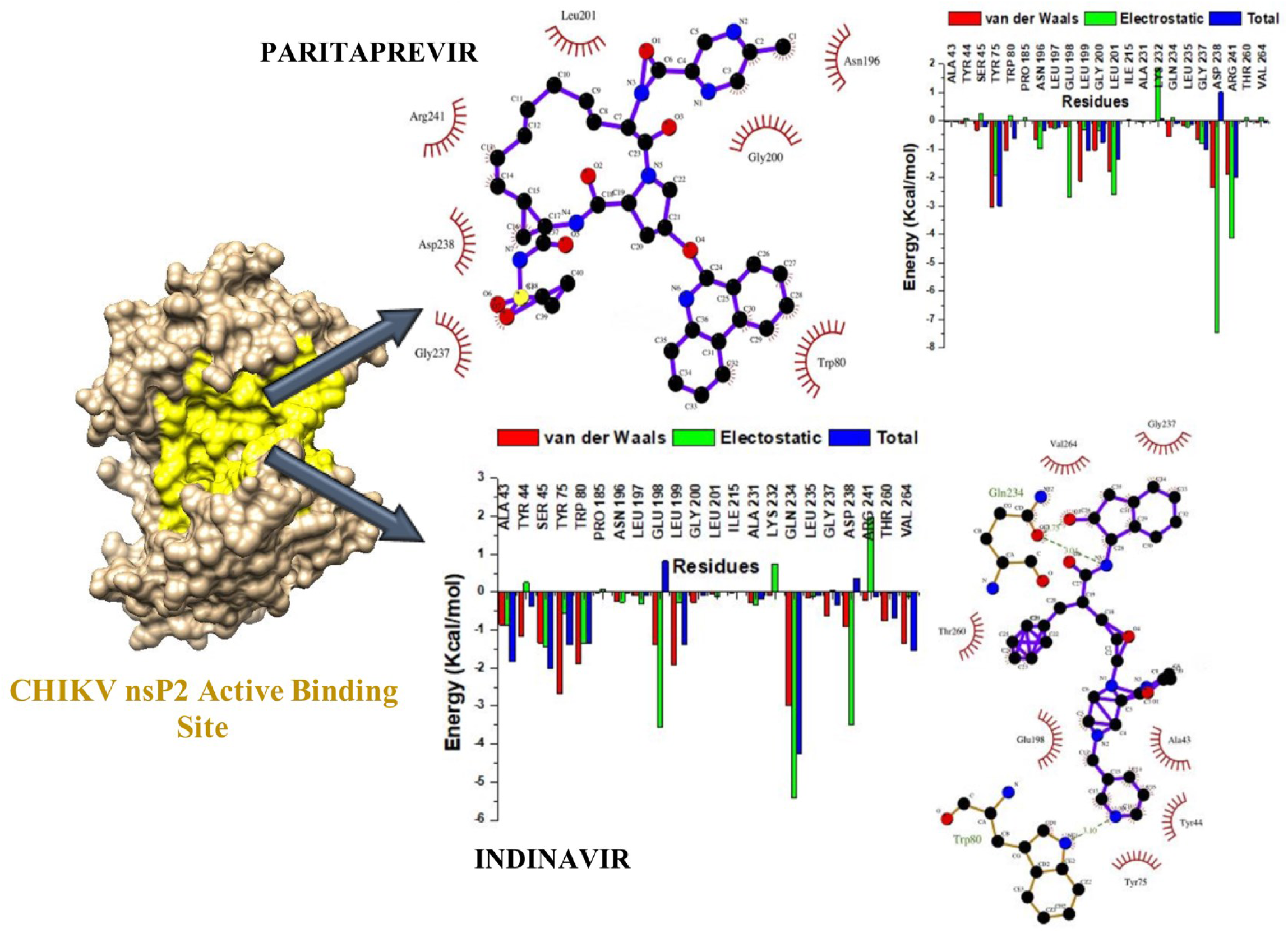
The per-residue free energy decomposition of Indinavir and Paritaprevir bound to CHIKV nsP2.

The radius of gyration lowers as the system becomes more compact. The results of Fig. 6 show that the compactness of the bound systems was lower than that of the apo system. The paritaprevir complex system (violet) showed a sharp decrease in compactness around 30 ns which was due to the slight movement of the ligand in the pocket (Kumar et al., 2019). This decrease in compactness advocates for the decrease in stability that was observed for the same system in RMSD. The overall compactness of the enzyme was lowered by the binding of ligands in the active site.

### 3.4 Principal Component Analysis

Principal component analysis (PCA) was used to characterize the conformational variations of the various ligands. PCA is a tool commonly used for conformational variations, they are used to understand the change of protein conformational diversities which are induced by the binding of various inhibitors (Halder et al., 2019). The conformational motions of all the systems were plotted on the two principal components (PC1 *vs* PC2) in order to gain a comprehensive understanding of the conformational changes in the systems.

The results of PCA show that the movements of the free enzyme atoms were more scattered than those of the bound systems. The movements of the atoms decrease from the complexes of Demethoxycurcumin to Grazoprevir and then Indinavir, with Paritaprevir showing the least movements. These results advocate for those of RMSF, where Paritaprevir decreased the flexibility of the enzyme the most. This means that Paritaprevir and Indinavir have reduced the activity of the protease the most via stable binding.

### 3.5 Binding Free Energy Calculations

As observed in Table 2, Indinavir, an HIV protease inhibitor, showed the highest binding free energy (*ΔG_bind_)* at -47.3135 kcal/mol. This was followed by HCV protease inhibitors, Grazoprevir and Paritaprevir, with binding free energies of -36.4226 kcal/mol and -31.3201 kcal/mol, respectively. The lowest binding free energy was exhibited by Demethoxycurcumin. The data provided in table 3 strongly suggests that Indinavir is one of the best inhibitors in comparison to the other inhibitors tested. Grazoprevir and Paritaprevir showed very favourable results as well considering that their binding free energies are higher than that of Demethoxycurcumin. The information in this table shows that van der Waals interactions were major contributors to the binding of the ligands in the active site. It should also be noted that all interactions were higher than those of Demethoxycurcumin for all the other ligands, except for electrostatic interactions for Grazoprevir. It should also be noted that the solvent, water, made no contributions towards the binding.

**Table 1:**
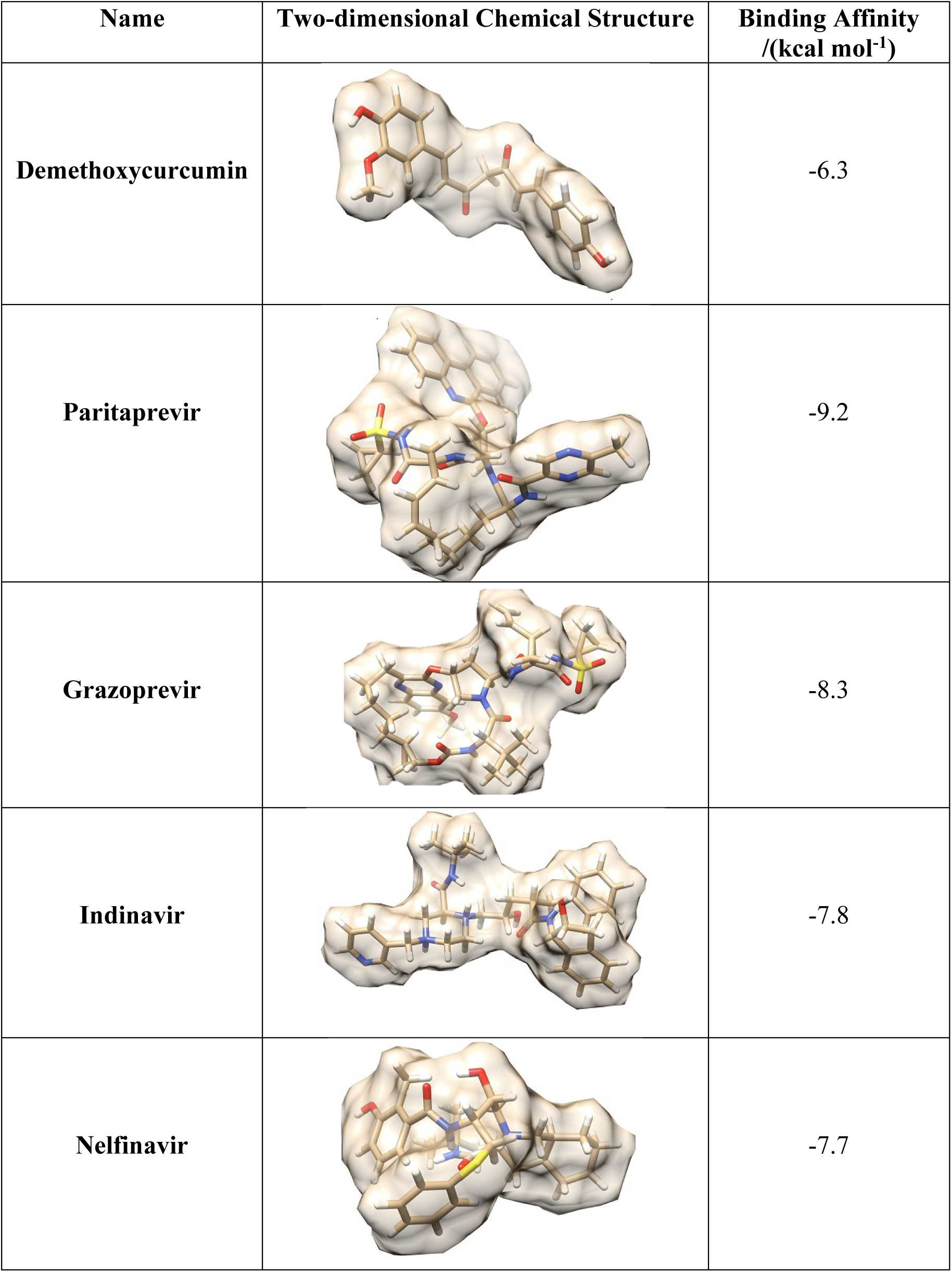
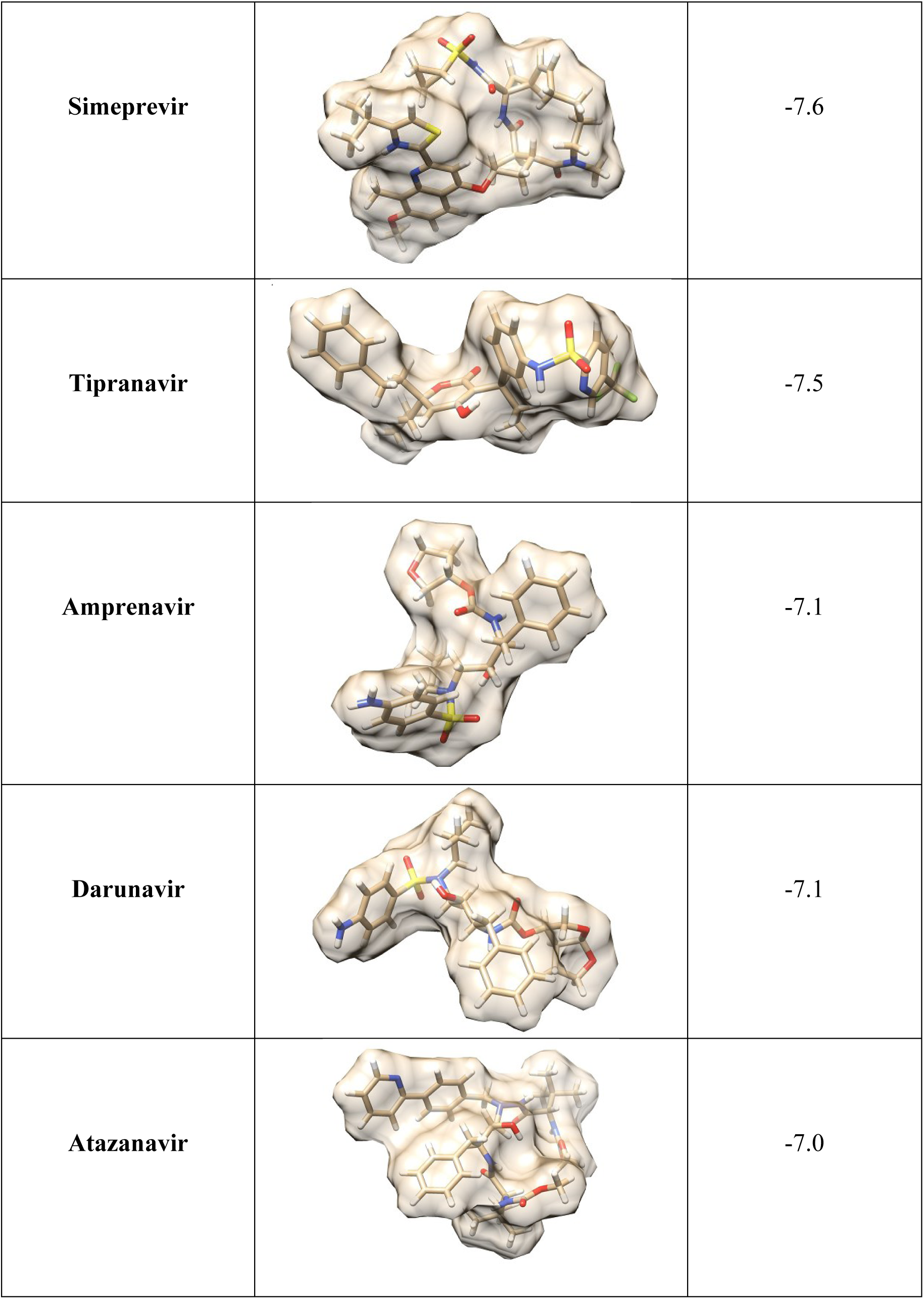
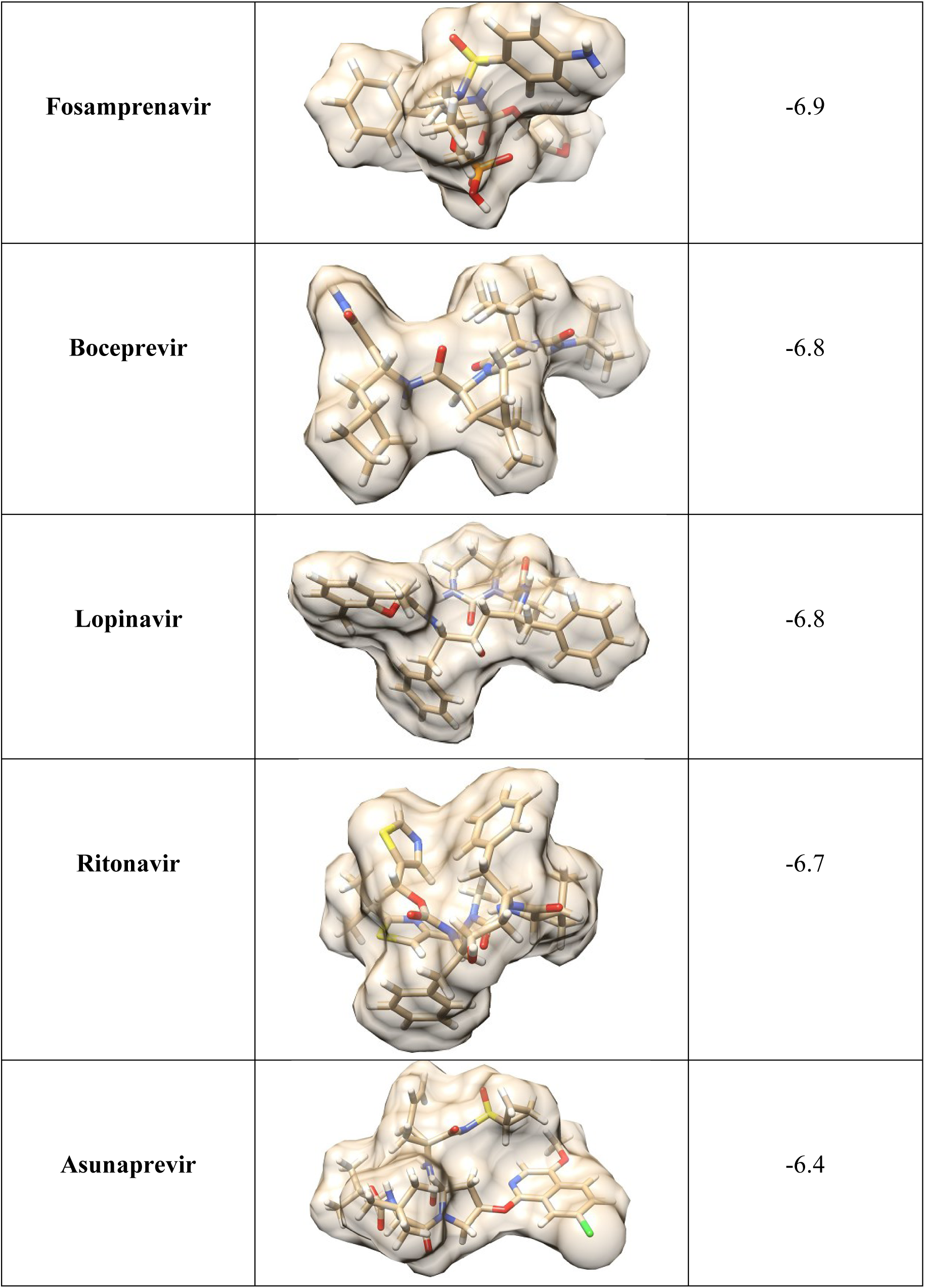
Structures and binding affinities of the chosen ligands for MD simulations.

| Name | Two-dimensional Chemical Structure | Binding Affinity<br>/(kcal mol<sup>-1</sup>) |
| --- | --- | --- |
| Demethoxycurcumin |  | -6.3 |
| Paritaprevir |  | -9.2 |
| Grazoprevir |  | -8.3 |
| Indinavir |  | -7.8 |
| Nelfinavir |  | -7.7 |
| **Simeprevir** |  | -7.6 |
| **Tipranavir** |  | -7.5 |
| **Amprenavir** |  | -7.1 |
| **Darunavir** |  | -7.1 |
| **Atazanavir** |  | -7.0 |
| **Fosamprenavir** |  | -6.9 |
| **Boceprevir** |  | -6.8 |
| **Lopinavir** |  | -6.8 |
| **Ritonavir** |  | -6.7 |
| **Asunaprevir** |  | -6.4 |

**Table 2:** MMPBSA based on Binding free energy of the complexes of Demethoxycurcumin, Grazoprevir, Indinavir and Paritaprevir with CHIKV nsp2.

|  | Energy Components (kcal/mol<sup>-1</sup>) |  |  |  |  |
| --- | --- | --- | --- | --- | --- |
| | $\Delta G_{bind}$ | $\Delta E_{ele}$ | $\Delta E_{vdw}$ | $\Delta E_{gas}$ | $\Delta G_{sol}$ |
| **Demethoxycurcumin** | -24.9816 | -28.3208 | -34.9907 | -63.3116 | 38.3300 |
| **Grazoprevir** | -36.4226 | -14.0681 | -62.3118 | -76.3799 | 39.9573 |
| **Indinavir** | -47.3135 | -35.5083 | -58.2327 | -93.7410 | 46.4275 |
| **Paritaprevir** | -31.3201 | -57.7499 | -50.7574 | -108.5073 | 77.1870 |

$\Delta E_{VDW}$ : van der Waal; $\Delta E_{elec}$ : electrostatic; $\Delta G_{gas}$ : non-polar solvation interaction; $\Delta G_{solv}$ : solvation free energy; $\Delta G_{bind}$ : calculated total free binding energy

As depicted in the per-residue energy contribution plots, the ligands interact with many of the residues of the binding pocket. The larger total residue energy contributions (|ΔG_binding_| > - 1) were observed for Indinavir. The residues that contributed the most energy to this complex were Ala43, Ser45, Tyr75, Trp80, Leu199, Gln234 and Val264. The highest energy contributors for this complex were van der Waals interactions of Tyr44 [-1.152 kcal/mol], Ser45 [-1.331 kcal/mol], Tyr75 [-2.673 kcal/mol], Trp80 [-1.881 kcal/mol], Glu198 [-1.377 kcal/mol], Leu199 [-1.919 kcal/mol], Gln234 [-2.986 kcal/mol] and Val264 [-1.351 kcal/mol]. The electrostatic interactions that were most favourable to the binding of this complex were from residues Ser45 [-1.430 kcal/mol], Trp80 [-1.332 kcal/mol], Gln234 [-5.418 kcal/mol], Asp238 [-3.505 kcal/mol] and Arg241 [-1.979 kcal/mol]. Indinavir formed hydrogen bonds with Gln234 and Trp80, while Demethoxycurcumin formed a hydrogen bond with Lys232. Van der Waals interactions were the highest contributors to the bonding interactions between all the ligands and the residues of the active site. Demethoxycurcumin was also found to interact more with the polar residues, while Grazoprevir, Indinavir and Paritaprevir interacted mostly with the non-polar residues. Trp80 is a substrate binding residue and forms part of a flexible loop that opens to the binding site. The formation of a hydrogen bond between Indinavir and Trp80 closes this flexible loop thus preventing the entrance into the active site and further bonding of other substrates is prohibited.

## 4 Conclusion

CHIKV is rapidly becoming a burden as the number of new cases increases thus enforcing the re-emergence status of the virus. A matter that has been substantially intensified by the evident lack of an adequately effective treatment or successful vaccine. Non-structural protease nsP2 protein plays a vital role in virus replication, therefore is a good target for the potential inhibition of CHIKV. In this study we use computational approaches to compare the inhibition abilities of Demethoxycurcumin to FDA approved HIV/HCV protease inhibitors; Grazoprevir, Indinavir and Paritaprevir. The results of the study show that the complexes of Indinavir and Paritaprevir are more stable and compact than the complex of Demethoxycurcumin. Indinavir, Grazoprevir and Paritaprevir also formed stable interactive bonds with conserved residues such as Cys10, Trp80 and Asp238. Notably, Indinavir formed a hydrogen bond interaction with Trp8,0 which shifted the conformation of the enzyme resuting in the closure of the flexible loop. Thus rendering the binding site inaccessible, ultimately leading to disruption of downstream essential activities of the nsP2 protein and impedement of viral replication. Based on the findings of this study, Indinavir should be further investigated for its inhibition potential using *in vitro* and thereafter *in vivo* methods. Furthermore, Grazoprevir, Indinavir and Paritaprevir could also be used as backbone molecules that require further optimisation and enhancements to improve binding and inhibition activity as potential therapeutic CHIKV agents.

## Acknowledgments

The authors would like to thank the National Research Foundation (NRF, South Africa), College of Health Sciences (CHS), University of KwaZulu-Natal for financial support and The Centre of High Performance Computing (CHPC, https://www.chpc.ac.za), Cape Town, RSA, for computational resources.

## References

1 J. Strauss and E. Strauss, Microbiol. Rev., 1994, 58, 491–562.

2 R. Sharma, P. Kesari, P. Kumar and S. Tomar, Virology, 2018, 515, 223–234.

3 F. J. Burt, W. Chen, J. J. Miner, D. J. Lenschow, A. Merits, E. Schnettler, A. Kohl, P. A. Rudd, A. Taylor, L. J. Herrero, A. Zaid, L. F. P. Ng and S. Mahalingam, Lancet Infect. Dis., 2017, 17, e107–e117.

4 H. Singh, R. Mudgal, M. Narwal, R. Kaur, V. Singh, A. Malik, M. Chaudhary and S. Tomar, Biochimie, 2018, 149, 51–61.

5 F. Burt, M. Rolph, N. Rulli, S. Mahalingam and M. Heise, Lancet (London, England), 2012, 379, 662–671.

6 Chikungunya virus | CDC, https://www.cdc.gov/chikungunya/index.html, (accessed 9 August 2021).

7 J. Jain, A. Kumari, P. Somvanshi, A. Grover, S. Pai and S. Sunil, F1000Research, DOI:10.12688/F1000RESEARCH.12301.2.

8 P. Nguyen, H. Yu and P. Keller, Interdiscip. Sci., 2018, 10, 515–524.

9 T. Ahola and A. Merits, Chikungunya Virus, 2016, 75.

10 B. Subudhi, S. Chattopadhyay, P. Mishra and A. Kumar, Viruses, DOI:10.3390/V10050235.

11 E. F. da Silva-Júnior, G. O. Leoncini, É. E. S. Rodrigues, T. M. Aquino and J. X. Araújo-Júnior, Bioorg. Med. Chem., 2017, 25, 4219–4244.

12 P. T. V. Nguyen, H. Yu and P. A. Keller, J. Mol. Graph. Model., 2015, 57, 1–8.

13 P. Kumar, D. Kumar and R. Giri, Pathog. (Basel, Switzerland), DOI:10.3390/PATHOGENS8030128.

14 Y.-S. Law, A. Utt, Y. B. Tan, J. Zheng, S. Wang, M. W. Chen, P. R. Griffin, A. Merits and D. Luo, Proc. Natl. Acad. Sci., 2019, 116, 9558–9567.

15 M. Narwal, H. Singh, S. Pratap, A. Malik, R. Kuhn, P. Kumar and S. Tomar, Int. J. Biol. Macromol., 2018, 116, 451–462.

16 Y. A. Karpe, P. P. Aher and K. S. Lole, PLoS One, 2011, 6, 22336.

17 R. Abdelnabi, S. N. Amrun, L. F. P. Ng, P. Leyssen, J. Neyts and L. Delang, Antiviral Res., 2017, 139, 79–87.

18 S. M. Corsello, J. A. Bittker, Z. Liu, J. Gould, P. McCarren, J. E. Hirschman, S. E. Johnston, A. Vrcic, B. Wong, M. Khan, J. Asiedu, R. Narayan, C. C. Mader, A. Subramanian and T. R. Golub, Nat. Med. 2017 234, 2017, 23, 405–408.

19 F. Pivari, A. Mingione, C. Brasacchio and L. Soldati, Nutrients, DOI:10.3390/NU11081837.

20 K. Mansouri, S. Rasoulpoor, A. Daneshkhah, S. Abolfathi, N. Salari, M. Mohammadi, S. Rasoulpoor and S. Shabani, BMC Cancer 2020 201, 2020, 20, 1–11.

21 S. B. Kutluay, J. Doroghazi, M. E. Roemer and S. J. Triezenberg, Virology, 2008, 373, 239.

22 V. H. Ferreira, A. Nazli, S. E. Dizzell, K. Mueller and C. Kaushic, PLoS One, 2015, 10, e0124903.

23 M. C. Fadus, C. Lau, J. Bikhchandani and H. T. Lynch, J. Tradit. Complement. Med., 2017, 7, 339–346.

24 B. Mounce, T. Cesaro, L. Carrau, T. Vallet and M. Vignuzzi, Antiviral Res., 2017, 142, 148–157.

25 S. Z. Moghadamtousi, H. A. Kadir, P. Hassandarvish, H. Tajik, S. Abubakar and K. Zandi, Biomed Res. Int., DOI:10.1155/2014/186864.

26 L. Padilla-S, A. Rodríguez, M. Gonzales, J. Gallego-G and J. Castaño-O, Arch. Virol., 2014, 159, 573–579.

27 A. Mazumder, K. Raghavan, J. Weinstein, K. Kohn and Y. Pommier, Biochem. Pharmacol., 1995, 49, 1165–1170.

28 D. Maksimovic-Ivanic, P. Fagone, J. McCubrey, K. Bendtzen, S. Mijatovi and F. Nicoletti, Int. J. cancer, 2017, 140, 1713–1726.

29 W. A. Chow, C. Jiang and M. Guan, Lancet Oncol., 2009, 10, 61–71.

30 M. D. Hanwell, D. E. Curtis, D. C. Lonie, T. Vandermeerschd, E. Zurek and G. R. Hutchison, J. Cheminform., 2012, 4, 17.

31 Z. Yang, K. Lasker, D. Schneidman-Duhovny, B. Webb, C. C. Huang, E. F. Pettersen, T. D. Goddard, E. C. Meng, A. Sali and T. E. Ferrin, J. Struct. Biol., 2012, 179, 269–278.

32 E. F. Pettersen, T. D. Goddard, C. C. Huang, G. S. Couch, D. M. Greenblatt, E. C. Meng and T. E. Ferrin, J. Comput. Chem., 2004, 25, 1605–1612.

33 O. Trott and A. J. Olson, J. Comput. Chem., 2010, 31, 455–61.

34 O. Trott and A. J. Olson, J. Comput. Chem., 2010, 31, 445–461.

35 A. W. Götz, M. J. Williamson, D. Xu, D. Poole, S. Le Grand and R. C. Walker, J. Chem. Theory Comput., 2012, 8, 1542–1555.

36 R. Salomon-Ferrer, D. A. Case and R. C. Walker, Wiley Interdiscip. Rev. Comput. Mol. Sci., 2013, 3, 198–210.

37 R. Salomon-Ferrer, A. W. Götz, D. Poole, S. Le Grand and R. C. Walker, J. Chem. Theory Comput., 2013, 9, 3878–3888.

38 L. Shunmugam and M. E. S. Soliman, RSC Adv., 2018, 8, 42210–42222.

39 N. M. Buthelezi, N. N. Mhlongo, D. G. Amoako, A. M. Somboro, S. C. Sosibo, L. Shunmugam, K. E. Machaba and H. M. Kumalo, 10.1080/07391102.2019.1677501 2019, 38, 4344–4352.

40 A. Bornot, C. Etchebest and A. de Brevern, Proteins, 2011, 79, 839–852.

41 M. Y. Lobanov, N. S. Bogatyreva and O. V. Galzitskaya, Mol. Biol. 2008 424, 2008, 42, 623–628.

42 I. Massova and P. A. Kollman, Perspect. Drug Discov. Des. 2000 181, 2000, 18, 113–135.

43 P. A. Kollman, I. Massova, C. Reyes, B. Kuhn, S. Huo, L. Chong, M. Lee, T. Lee, Y. Duan, W. Wang, O. Donini, P. Cieplak, J. Srinivasan, D. A. Case and T. E. Cheatham, Acc. Chem. Res., 2000, 33, 889–897.

44 A. K. Halder and B. Honarparvar, Struct. Chem., 2019, 30, 1715–1727.

